# Multiscale functional connectivity patterns of the aging brain learned from rsfMRI data of 4,259 individuals of the multi-cohort iSTAGING study

**DOI:** 10.1101/2022.07.27.501626

**Authors:** Zhen Zhou, Dhivya Srinivasan, Hongming Li, Ahmed Abdulkadir, Ilya Nasrallah, Junhao Wen, Jimit Doshi, Guray Erus, Elizabeth Mamourian, Nick R. Bryan, David A. Wolk, Lori Beason-Held, Susan M. Resnick, Theodore D. Satterthwaite, Christos Davatzikos, Haochang Shou, Yong Fan, the ISTAGING Consortium

## Abstract

To learn multiscale functional connectivity patterns of the aging brain, we built a brain age prediction model of functional connectivity measures at seven scales on a large fMRI dataset, consisting of resting-state fMRI scans of 4259 individuals with a wide age range (22 to 97 years, with an average of 63) from five cohorts. We computed multiscale functional connectivity measures of individual subjects using a personalized functional network computational method, harmonized the functional connectivity measures of subjects from multiple datasets in order to build a functional brain age model, and finally evaluated how functional brain age gap correlated with cognitive measures of individual subjects. Our study has revealed that functional connectivity measures at multiple scales were more informative than those at any single scale for the brain age prediction, the data harmonization significantly improved the brain age prediction performance, and harmonization in the tangent space worked better than in the original space. Moreover, brain age gap scores of individual subjects derived from the brain age prediction model were significantly correlated with clinical and cognitive measures. Overall, these results demonstrated that multiscale functional connectivity patterns learned from a large-scale multi-site rsfMRI dataset were informative for characterizing the aging brain and the derived brain age gap was associated with cognitive and clinical measures.

## Introduction

Brain age derived from non-invasive magnetic resonance imaging (MRI) data using machine learning provides a novel means to quantify brain development and aging process (Douaud et al., 2014; Cole and Franke, 2017). Brain age gap (BAG), quantifying the difference between the brain age and the chronological age, has demonstrated promising performance for elucidating atypical brain development and aging (Cole and Franke, 2017; Truelove-Hill et al., 2020).

Most brain age modeling studies focused on structural MRI data and have shown that brain age is associated with changes in both gray matter (GM) (Erus et al., 2015; Chung et al., 2017; Minkova et al., 2017; Truelove-Hill et al., 2020) and white matter (WM) (Prins and Scheltens, 2015; Habes et al., 2016; Habes et al., 2021). Particularly, it has been demonstrated that the brain age derived from structural MRI data can accurately delineate trajectories of brain development and identify individuals with cognitive precocity or delay (Erus et al., 2015). Similarly, it has been shown that brain development during adolescence is associated with widespread, regionally hierarchical gray matter loss and white matter increase (Truelove-Hill et al., 2020). A recent brain age modeling study also revealed brain aging trajectories in a large cohort (Habes et al., 2021). Moreover, large-scale neuroimaging studies have revealed that brain disorders are associated with brain age gap estimated from structural neuroimaging data (Kaufmann et al., 2019; Bashyam et al., 2020; Dinsdale et al., 2021).

The brain age has also been investigated based on functional neuroimaging data (Dosenbach et al., 2010; Dennis and Thompson, 2014; Li et al., 2018; Zonneveld et al., 2019; Truelove-Hill et al., 2020). It has been demonstrated that brain maturity can be accurately estimated based on individual subjects’ functional connectivity (FC) measures computed from functional MRI (fMRI) data (Dosenbach et al., 2010). Multiple studies have reported that the brain age is associated with changes in widespread functional network connectivity measures over the course of adolescence (Fair et al., 2008; Di Martino et al., 2014; Truelove-Hill et al., 2020). A large population-based aging study has revealed that brain aging is associated with weak FC within the anterior default mode network (DMN), ventral/salience attention network (VAN), and somatomotor network (SMN) and strong FC within the visual network (VN) (Zonneveld et al., 2019). However, most existing functional neuroimaging-based brain aging studies typically focused on data from single datasets (Dosenbach et al., 2010; Chan et al., 2014; Liang et al., 2019; Zonneveld et al., 2019), lacking diversity in the study cohorts.

In this study, we investigated functional connectivity patterns of the aging brain based on fMRI scans of a diverse cohort (n = 4259) from five different sites in a brain age modeling framework. We computed functional connectivity measures of individual subjects at multiple scales using a personalized functional network computational method (Li et al., 2017), harmonized the multiscale functional connectivity measures of different sites in their tangent space using ComBat-GAM (Pomponio et al., 2020), built a regression model on the harmonized functional connectivity measures to estimate the brain age and characterize the brain age gap, and finally we identified clinical and cognitive measures that were significantly correlated with the brain age gap in order to investigate whether the functional brain age gap is associated with cognitive functions and biological measures.

## Materials and methods

The overall design of this study for characterizing multiscale FC patterns of the aging brain is schematically illustrated in Figure 1, including computing multiscale FC measures of individual subjects, harmonizing the FC measures of multiple datasets, building a brain age prediction model to identify informative FC measures, and finally evaluating how the brain age gap correlated with cognitive/biological measures of individual subjects.

**Figure 1.**
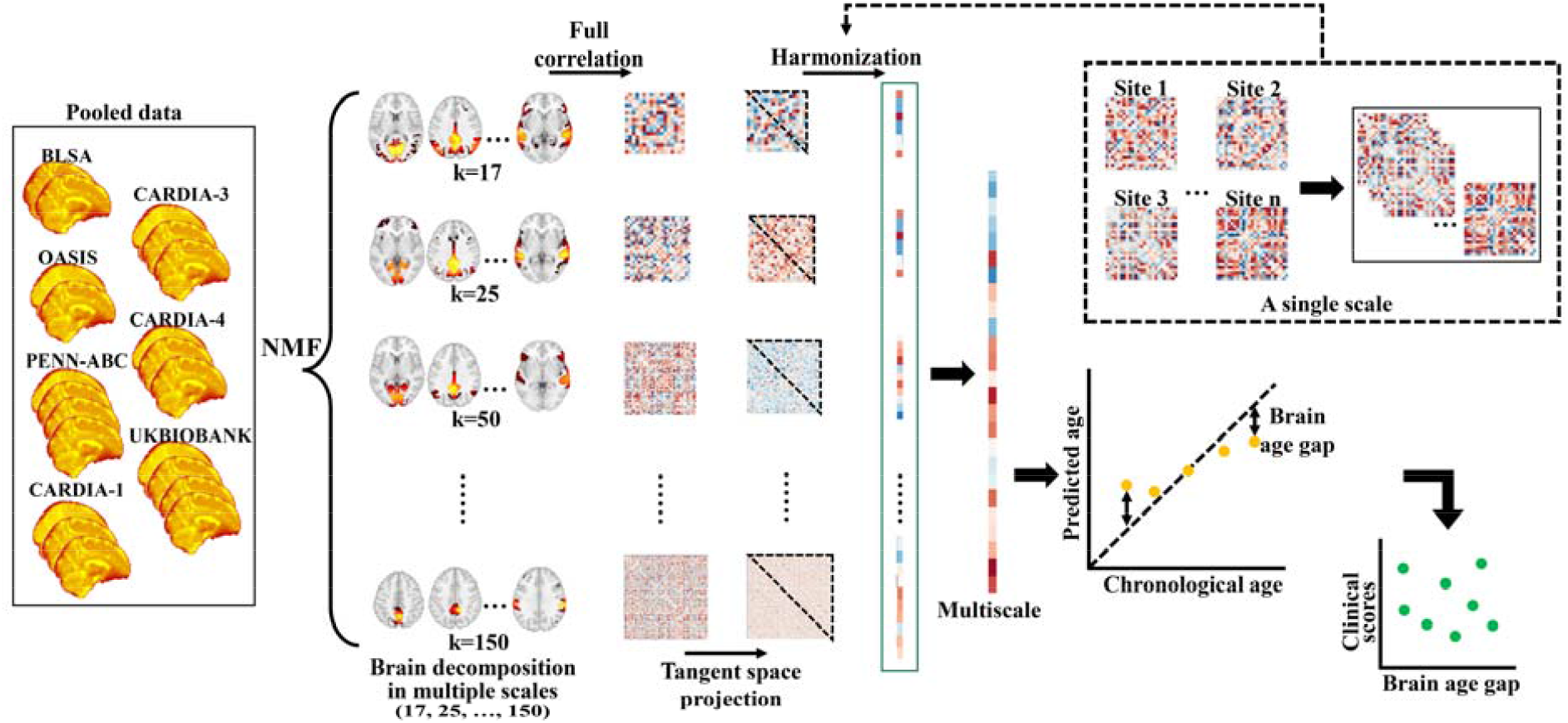
An overall flowchart of the present study. Multiscale functional connectivity measures were computed using a personalized functional network computing method, projected into their tangent space, harmonized using a Combat method, and finally used as input to build a brain age prediction model, from which a brain age gap score was derived to charactering the aging brain based functional MRI data of individuals.

### Participants

We identified 4549 participants with resting-state fMRI data from the iSTAGING (Imaging-based coordinate SysTem for AGIng and NeurodeGenerative diseases) consortium (Habes et al., 2021). The fMRI scans were collected from subjects at a wide age range (22 to 97 years) from 5 different cohorts, including the Baltimore Longitudinal Study of Aging (BLSA), the Open Access Series of Imaging Studies (OASIS-3), the Coronary Artery Risk Development in Young Adults (CARDIA), the University of Pennsylvania Aging Brain Cohort (PENN_ABC), and the UK (United Kingdom) Biobank. Particularly, the CARDIA cohort was divided into three subcohorts, namely CARDIA-1, CARDIA-3, and CARDIA-4, according to their scanners/sites used for the data collection. Since identical scanners and protocols were used for brain imaging scanning in the UKBiobank (Focke et al., 2011; Chen et al., 2014; Alfaro-Almagro et al., 2018), the UKBiobank scans were considered from one single data site. In summary, the fMRI scans were modeled from seven different sites, their demographic information and scanning protocols are summarized in Table 1. This study was approved by the supervisory committee and the institutional review board of the University of Pennsylvania School of Medicine.

**Table 1.**
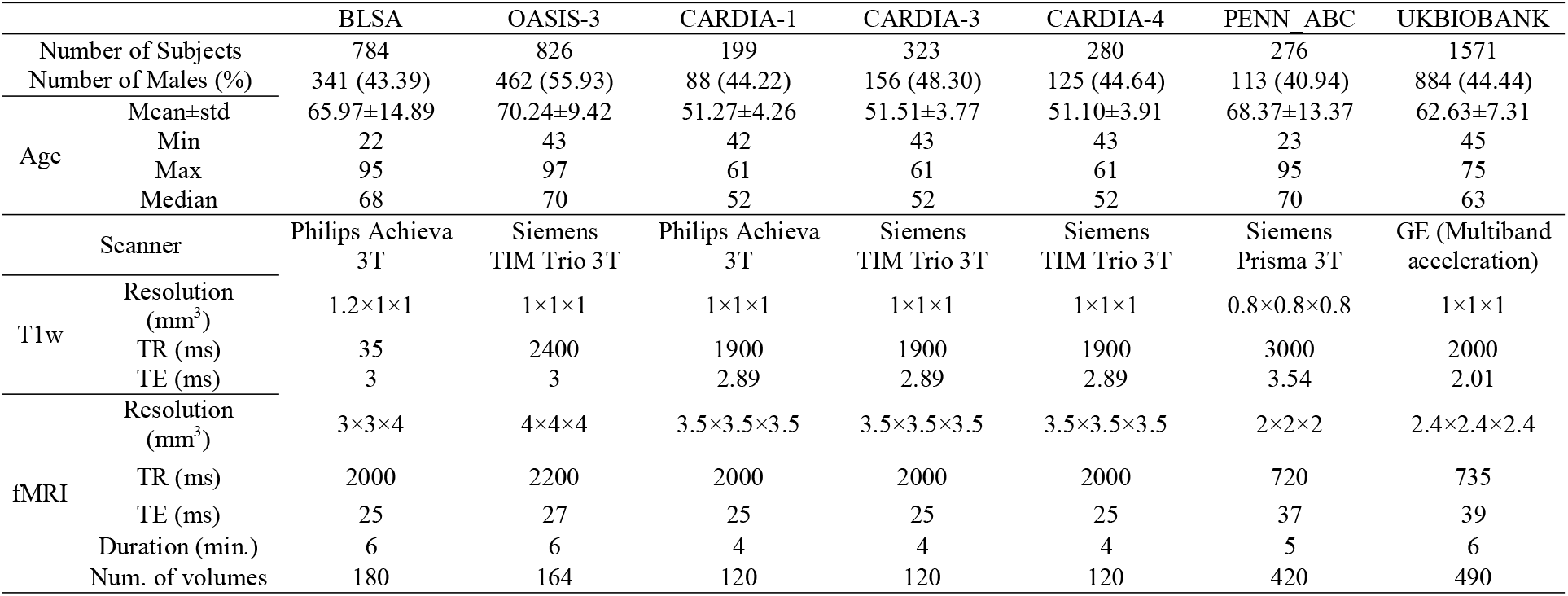
Demographic information and the scanning protocols of the multisite data

### Data processing

The UKBiobank scans were processed using UKBiobank preprocessing pipeline (Alfaro-Almagro et al., 2018), and other scans were processed using a modified UKBiobank preprocessing pipeline with steps, including head motion correction by FSL’s MCFLIRT (Jenkinson et al., 2012), global 4D mean intensity normalization, and temporal high-pass filtering (> 0.01 Hz). After these standard pre-processing steps, random noise was removed using FIX (FMRIB’s Independent Component Analysis-based Xnoiseifier) (Griffanti et al., 2014; Salimi-Khorshidi et al., 2014). Specifically, the FIX model was built upon WhII_Standard.RData from Whitehall Imaging Study (Filippini et al., 2014) due to its similarity with our rsfMRI data. The preprocessed rsfMRI scans were co-registered to their corresponding T1-weighted images using FLIRT with BBR as the cost function, and the T1-weighted images were registered to the MNI152 template using FSL’s FNIRT (non-linear registration), generating rsfMRI scans with a spatial resolution of 2 × 2 × 2 mm^3^. A brain mask was applied in standard space to exclude white matter, cerebral spinal fluid, and cerebellum (cerebellum are not fully covered for some subjects).

Participants were excluded from subsequent analyses if their rsfMRI scans had mean relative displacement higher than 0.2 mm, more than 60% of frames with motion exceeding 0.3 mm, or temporal signal-to-noise ratio (tSNR) smaller than 100 except for the UKBiobank scans that were acquired using a multiband protocol (Alfaro-Almagro et al., 2018). In total, 290 participants were excluded, and the remaining 4259 subjects were included in the following analyses.

### Computation of multiscale functional connectivity measures

Multiscale functional connectivity measures were computed from each preprocessed rsfMRI scan based on functional networks (FNs) obtained using a personalized functional network computational method (Li et al., 2017; Cui et al., 2020). We computed personalized FNs for each individual subject using a group-sparsity regularized non-negative matrix factorization (NMF) method (Li et al., 2017; Cui et al., 2020), which has been successfully adopted in multiple recent studies for computing personalized FNs (Cui et al., 2022; Pines et al., 2022; Shanmugan et al., 2022). Particularly, we first computed group-level FNs using a normalized-cuts based spectral clustering method to identify representative FNs from 50 sets of group-level FNs, each set being computed on a subset of 150 subjects randomly selected from each of the sites with a probability proportional to the sample sizes of different sites. The group-level FNs were then used as initializing FNs to compute personalized FNs based on each subject’s fMRI data.

Given a group of *n* subjects, each with fMRI data *X*^*i*^ ∈ *R*^*T*×*S*^, *i* = 1, …,*n* consisting of *S* voxels and *T* time points, we aim to find *K* non-negative FNs 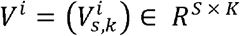 and their corresponding time series 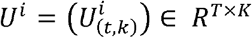 for each subject, such that

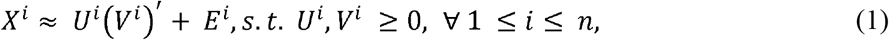

where (*V*^*i*^)′ is the transpose of *V*^*i*^, and *E*^*i*^ is additional independent noise following a Gaussian distribution. Both *U*^*i*^ and *V*^*i*^ are constrained to be non-negative so that each FN does not contain any anti-correlated functional units. A group consensus regularization term was applied to ensure inter-individual correspondence, which was implemented as a scale-invariant group sparsity term on each column of *V*^*i*^,*i* =1,…,*n* and formulated as

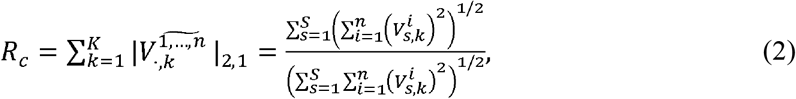

The data locality regularization term was applied to encourage spatial smoothness and coherence of the FNs using graph regularization and formulated as:

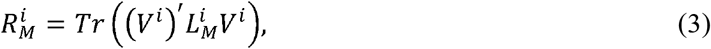

where 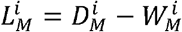 is a Laplacian matrix for subject 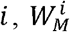 is a pairwise affinity matrix to measure spatial closeness or functional similarity between different voxels, and 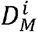 denotes its corresponding degree matrix. The affinity between each pair of spatially connected voxels is calculated as 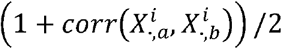, where 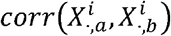 is the Pearson correlation coefficient between time series 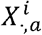 and 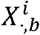 while others were set to be zero so that 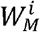 would be sparse. Finally, we identified subject specific functional networks by optimizing a joint model with integrated data fitting and regularization terms formulated as:

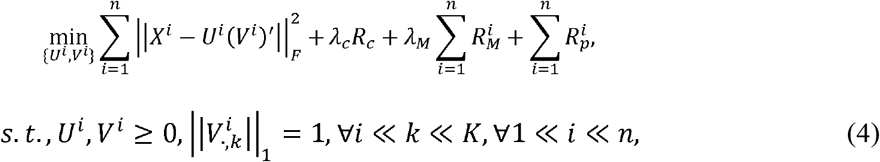

where λ_*M*_ and λ_*c*_ are used to balance the data fitting, data locality, and group consensus regularization terms with values setting as the those validated in the previous study (Li et al., 2017). Instead of computing the FNs at a specific spatial scale, we computed the FNs at seven scales, yielding seven sets of *K* (*K*=17, 25, 50, 75, 100, 125 and 150) FNs.

To facilitate the interpretation of the personalized FNs, we labeled each of them based on spatial overlapping between their group-level FNs and Yeo’s 7-network atlas, including the visual network (VN), somatomotor network (SMN), dorsal attention network (DAN), salience/ventral attention network (VAN), limbic network (LN), frontoparietal network (FPN) and default mode network (DMN).

### Sanity testing for computed personalized FNs

Quality control was carried out to ensure that the personalized FNs had higher functional homogeneity than their group-level counterparts and maintained good spatial correspondence with their group-level counterparts. Particularly, a FN’s functional homogeneity was measured by a weighted mean of the correlation coefficients between the time courses of all the voxels within the FN and its centroid time course that was calculated as a weighted mean time course over the FN with its voxel-wise loadings as weights. The spatial correspondence between the personalized FNs and the group-level FNs was evaluated based on pairwise spatial correlation coefficients. Specifically, each personalized FN is deemed to maintain correspondence with its corresponding group level counterpart if 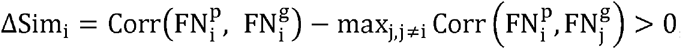, where 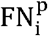 denotes the i-th personalized 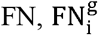 denotes its corresponding group average 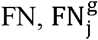 represents other group average FNs, and Corr(·,·) is the spatial correlation coefficient between two FNs.

### Harmonization of functional connectivity measures in tangent space

In order to alleviate site effect of functional connectivity measures of participants from different sites, we adopted ComBat-GAM to harmonize the functional connectivity measures (Pomponio et al., 2020). Since the functional connectivity measures themselves essentially resided on a Riemannian manifold (You and Park, 2021), we applied ComBat-GAM to the functional connectivity measures in their tangent space (Pervaiz et al., 2020). Functional connectivity measure between each pair of FNs within each set was estimated as Pearson correlation between time series of the FNs (Zhou et al., 2020), yielding seven sets of functional connectivity matrices ***C*** ∈ *R*^*K*×*K*^, *K*=17, 25, 50, 75, 100, 125 and 150. The tangent space projection of the FC matrix ***C*** was obtained through

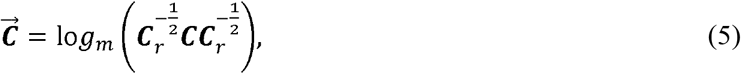

where ***C***_***r***_ ∈ *R*^*K*×*K*^ is a reference point in the manifold, *K* is the number FNs, lo*g*_*m*_ represents the logarithm operation on the FC matrix and 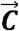 is the resulting FC measures in the tangent space. To ensure that all projected covariance matrices lie in the same tangent plane, we chose the geometric mean as the reference point (Fletcher et al., 2004; Ng et al., 2014; Yger et al., 2017). The FC measures 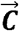 in the tangent space can be projected back to the original space by

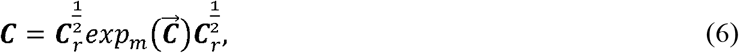

where *exp*_*m*_ is the exponential operation on the FC matrix.

We harmonized the FC measures in the tangent space using ComBat-GAM (Pomponio et al., 2020) with sex and age as covariates. After data harmonization, we vectorized the upper triangular part of the harmonized connectivity matrix in each scale and stacked them across scales to obtain a panel of multiscale features as illustrated in Figure 1.

### Experimental design and statistical analysis

#### Brain age predictive modeling

We built a brain age prediction model on the harmonized FC measures using Ridge regression by optimizing

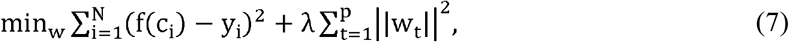

where *y*_*i*_ and *c*_*i*_ denote age and features of subject *i, p* denotes the number of features, *w*_*t*_ is a regression coefficient, *t* = 1,2, …,*p*, and λ is a regularization parameter. We adopted a nested 5-fold cross-validation, with the inner folds determining the optimal parameter λ within the grid of [2^3^,2^3.5^, 2^4^, …,2^7.5^, 2^8^] and the outer folds estimating the generalizability of the model. The cross-validation folds were generated randomly with comparable age distributions. The brain age prediction performance was quantified with mean absolute error (MAE) and correlation coefficient between the chronological age and the predicted age. The optimal λ value was determined based on MAE.

We also built brain age prediction models on FC measures in their original space, with and without the data harmonization, as well as FC measures in the tangent space without the data harmonization, respectively. In addition to the brain age prediction models built on the multiscale functional connectivity measures, we also built brain age prediction models on FC measures of individual scales with 17, 25, 50, 75, 100, 125 and 150 FNs, respectively. All the brain age prediction models were built and evaluated with the same nested 5-fold cross-validation. The performance difference between the prediction models built on FC measures with and without the data harmonization was assessed with Wilcoxon signed-rank test across individual scales and their combination.

#### Functional network connectivity measures informative for brain age prediction

A permutation test was performed to evaluate the statistical significance of individual FC measures and the accuracy of the brain age prediction model (Mourao-Miranda et al., 2005; Cui et al., 2018). Particularly, we permuted the age labels of all subjects and repeated the whole cross-validation process (splitting the whole dataset into training and testing subsets) with the optimal λ parameter for 1000 times, yielding 1000 null brain age prediction models. We projected the weight derived from the model back to the original space for interpretation with Equation (6) using the same reference matrix. Thus, we obtained 1000 weight vectors and projected them back to original space. We also projected the feature weight from the real model without permuting the labels back to original space. The *p* value for each feature is the proportion of permutations that showed a higher weight value than the actual value from the real model. And those features with *p* < 0.05 were identified as the significantly contributing network connectivity features. Each model’s regression coefficients were projected back to the original space of their associated FC measures for quantifying their contribution to the brain age prediction. Furthermore, correlation between individual FC measures and the chronological age was also calculated to identify FC measures significantly associated with aging.

#### Exploration of association between the brain age gap and clinical/cognitive score

Cognitive and clinical measures available for the majority of the 4259 subjects were curated to investigate if they are correlated with the brain age gap. We computed correlations between the BAG and cognitive and physiological functioning measures of individual subjects. As summarized in Table 3, the cognitive and physiological functioning measures included attention, executive function, working memory, and verbal, spanning several cognitive domains, as well as Systole, Diastole, and BMI (Body Mass Index). Pearson’s correlations between the BAG and these measures were computed with sex, age, and site as covariates (Dinsdale et al., 2021). The BAG was computed with the brain age prediction bias corrected using a linear regression method (Beheshti et al., 2019).

## Results

### Sanity testing results

We performed the quality control of personalized FNs at all different spatial scales (17, 25, …, 150), and found that all the personalized FNs had higher functional homogeneity than their corresponding group-level FNs and they maintained good correspondence with their corresponding group-level FNs, i.e., ΔSim_i_ was larger than 0 for each personalized FN. Overall, the generated personalized brain functional networks provided an improved fit to each individual’s fMRI data than to the group-level FNs that are not equipped to characterize inter-individual variations in functional neuroanatomy.

### Brain age prediction results

The prediction performance of all the brain age prediction models under comparison is summarized in Table 2 and illustrated in Figure 2, indicating that the brain age prediction model built on the harmonized multiscale FC features in the tangent space obtained the best age prediction performance, with a mean MAE of 5.57 years and a mean correlation coefficient of 0.78. The permutation test indicated that its prediction performance was statistically significant with *p* < 0.001 in terms of both MAE and correlation coefficient. The data harmonization did improve the performance if the models were built on FC measures in the tangent space (*p* = 0.025 for MAE and *p* = 0.006 for correlation coefficient, Wilcoxon signed-rank test across individual scales and their combinations) but did not consistently improve the models built on FC measures in their original space.

**Table 2.** Brain age prediction performance (MAE and correlation between the predicted and chronological ages) of all brain age prediction models built on FC measures of individual scales and their combination with and without the data harmonization (H+ and H-denote the models built on the FC measures harmonized or not, respectively)

| Scales | MAE (year) between<br>predicted age and chronological age |  |  |  | Correlation coefficient between<br>predicted age and chronological age |  |  |  |
| --- | --- | --- | --- | --- | --- | --- | --- | --- |
|  | Original Space |  | Tangent Space |  | Original Space |  | Tangent Space |  |
|  | H- | H+ | H- | H+ | H- | H+ | H- | H+ |
| 17 | 8.45±0.12 | 8.34±0.14 | 8.01±0.11 | 7.96±0.12 | 0.42±0.03 | 0.42±0.03 | 0.49±0.03 | 0.51±0.03 |
| 25 | 7.88±0.16 | 8.12±0.17 | 7.38±0.15 | 7.43±0.18 | 0.52±0.03 | 0.47±0.03 | 0.58±0.03 | 0.58±0.02 |
| 50 | 7.67±0.14 | 8.23±0.12 | 7.21±0.16 | 7.33±0.16 | 0.56±0.03 | 0.46±0.02 | 0.64±0.01 | 0.62±0.02 |
| 75 | 7.92±0.18 | 8.74±0.27 | 7.91±0.11 | 8.17±0.19 | 0.55±0.02 | 0.41±0.03 | 0.61±0.08 | 0.59±0.01 |
| 100 | 8.28±0.13 | 9.30±0.16 | 8.02±0.08 | 7.80±0.10 | 0.53±0.02 | 0.36±0.02 | 0.60±0.07 | 0.61±0.01 |
| 125 | 8.26±0.24 | 8.88±0.25 | 7.19±0.12 | 6.96±0.15 | 0.54±0.03 | 0.42±0.04 | 0.65±0.02 | 0.67±0.02 |
| 150 | 8.56±0.19 | 8.86±0.09 | 6.70±0.09 | 6.45±0.09 | 0.52±0.04 | 0.42±0.03 | 0.68±0.02 | 0.71±0.02 |
| Multiscale | 7.65±0.17 | 8.29±0.18 | 5.92±0.11 | 5.57±0.11 | 0.59±0.03 | 0.46±0.03 | 0.76±0.02 | 0.78±0.01 |

**Figure 2.**
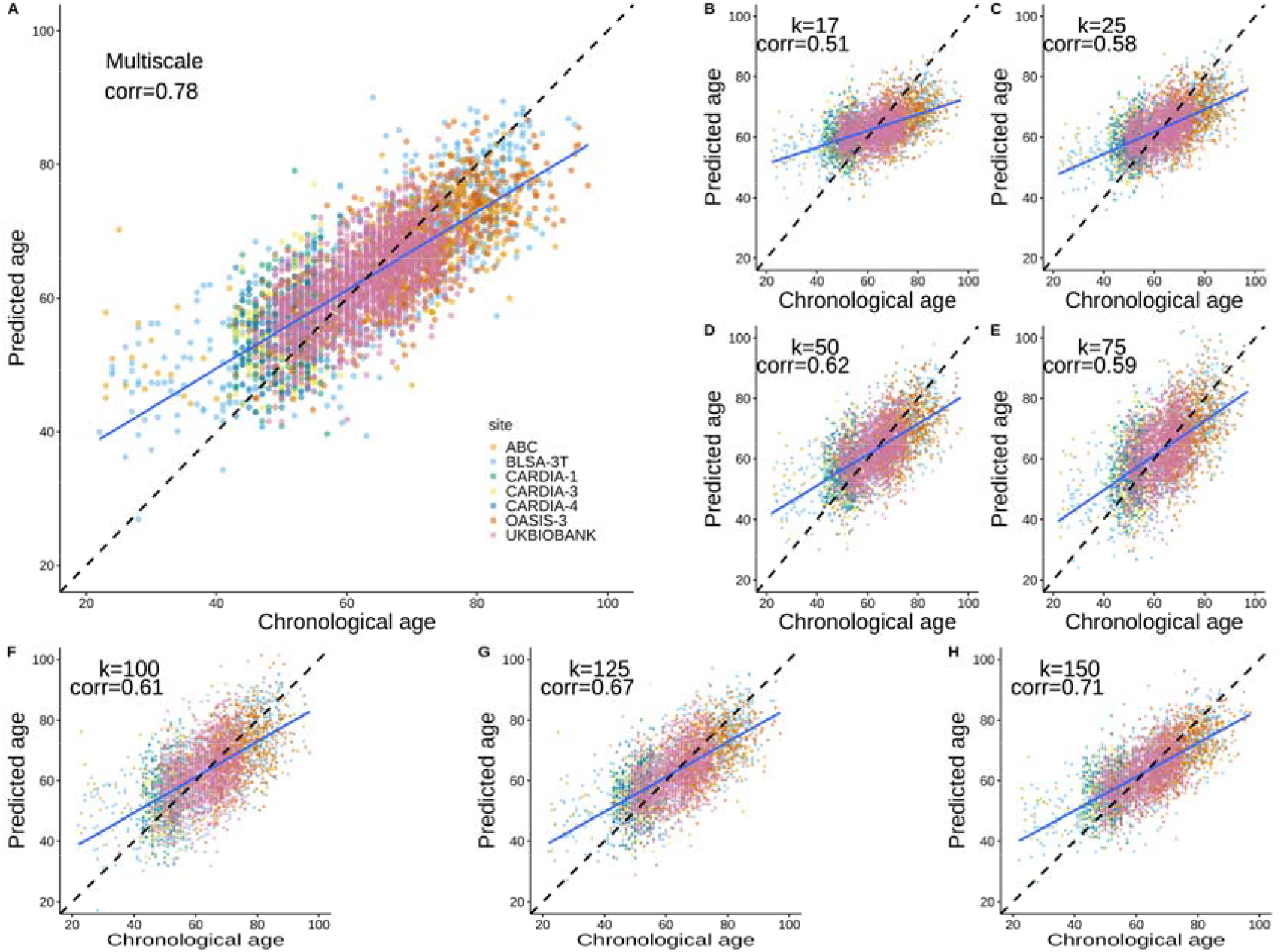
Correlations between the chronological age and the predicted age obtained by prediction models built upon functional connectivity measures of individual scales and their combination. In each subplot, x-axis represents the chronological age while y-axis denotes the predicted age. (A): brain age was predicted by the prediction model built upon the functional connectivity measures of all individual spatial scales; (B)-(F): brain age was predicted by the prediction model built upon the functional connectivity measures between functional networks computed with the settings of 17, 25, 50, 75, 100, 125, and 150 FNs separately. In each subplot, the black dashed line is the identity line while the blue solid line indicates the best linear fit of the predicted age to the chronological age.

Figure 2 shows scatter plots and associated correlation coefficients between the predicted and chronological ages obtained by the brain age prediction models built on the harmonized FC measures of individual scales and their combination in the tangent space, indicating again that the brain age prediction model built on the harmonized multiscale FC features in the tangent space obtained the best age prediction performance.

### Informative functional connectivity measures for predicting brain age

Figure 3 shows a hierarchical organization of the FNs from coarse to finer scales and FC measures contributed to the brain age prediction model built on the harmonized multiscale FC features in the tangent space with statistical significance (43 FC measures in total, *p* < 0.05, permutation test), illustrating that FC measures at multiple scales were informative for the brain age prediction. Particularly, FC measures between DMN, frontoparietal network (FPN), VN, VAN, SMN, dorsal attention network (DAN) and limbic network (LN) played vital roles. Among 43 informative FC measures identified by the brain age prediction model, 19 were significantly correlated with aging (*p<0*.*05*) and most of them were connections between SMN, VAN, DMN or FPN, as illustrated in Figure 4.

**Figure 3.**
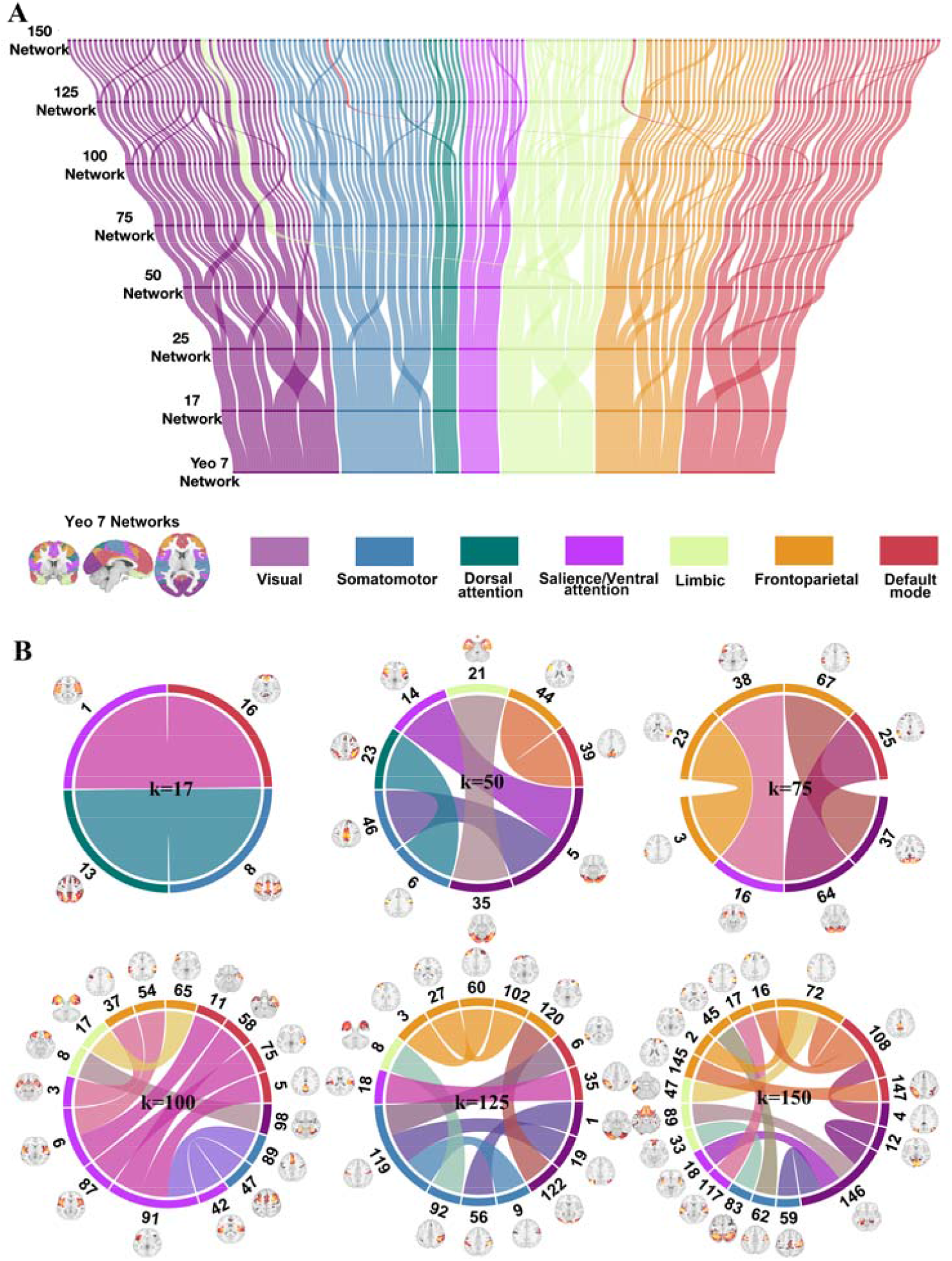
Functional connections informative for predicting the brain age. (A): A multi-scale organization of the brain networks, illustrated following the Yeo atlas of 7 networks, including VN, SMN, DAN, VAN, LN, FPN and DMN. (B): Functional connections informative for the brain age prediction. The different edge colors are used to differentiate different brain networks, corresponding to the FNs of the Yeo’s 7-network parcellation.

**Figure 4.**
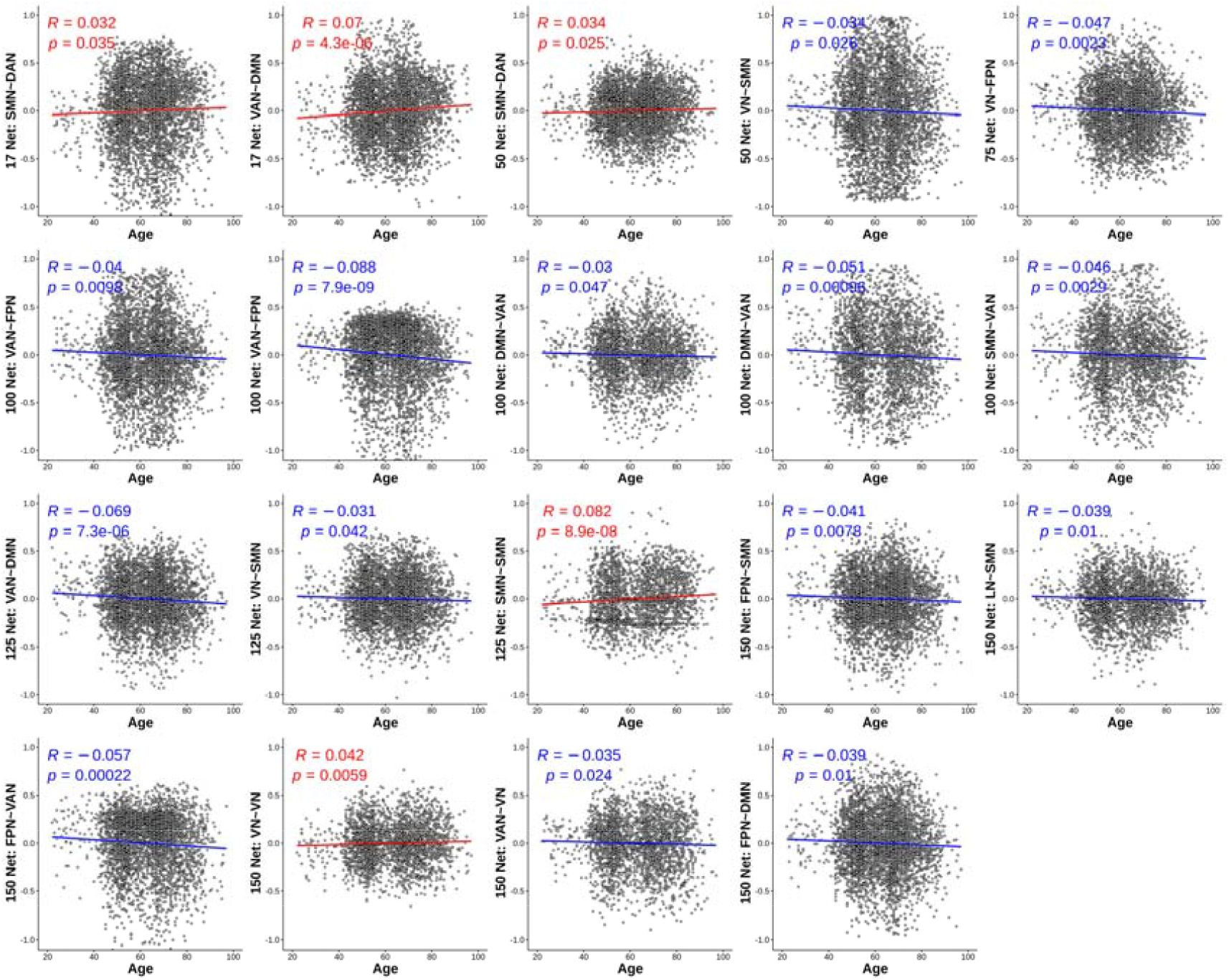
Functional connections with connectivity significantly correlated with the chronological. Regression lines in red and blue indicate functional connectivity measures positively and negatively correlated with the chronological age, respectively.

### Brain age gap correlation with cognitive scores

Table 3 summarizes correlation coefficients between BAG and clinical/cognitive measures. Particularly, MMSE (Mini-Mental State Examination), DSST (Digital Symbol Substitution Test), CVLT_IM (California Verbal Learning Test Immediate), CVLT_Long (California Verbal Learning Test Long), DSB (Digit Span Backward), and DSF (Digit Span Forward) were negatively correlated with the BAG, while measures of ‘time taken to complete a cognitive task’, i.e., TMT_A (Trail Making Test Part A) and TMT_B (Trail Making Test Part B) were correlated positively with the BAG. The BAG was positively correlated with blood pressure (systole and diastole) and BMI (Body Mass Index).

**Table 3.** Correlation between BAG and cognitive/clinical measures

|  | Clinical or cognitive measures | correlation coefficients | *p* values |
| --- | --- | --- | --- |
| Attention | DSF | -0.061 | 0.0109 |
|  | TMT_A | 0.078 | **0.0012\*** |
| Executive | DSST | -0.0957 | **0.0002\*** |
|  | TMT_B | 0.0641 | 0.0089 |
|  | DSB | -0.0779 | 0.0138 |
| Verbal Fluency | BNT | -0.1063 | 0.24 |
|  | ANI_Fluency | -0.1158 | 0.0664 |
|  | Category_Fluency | -0.0866 | 0.0163 |
|  | Letter_Fluency | -0.0667 | 0.0646 |
|  | VEG_Fluency | -0.1405 | 0.0137 |
| Verbal Memory | CVLT_IM | -0.0849 | 0.019 |
|  | CVLT_Long | -0.0919 | 0.0113 |
| Global | MMSE | -0.1362 | **0.0000\*** |
|  | MOCA | -0.146 | 0.0035 |
|  | WM_SUM | -0.1629 | 0.0171 |
| Biological | Systole | 0.0544 | 0.0032 |
|  | Diastole | 0.0148 | 0.4213 |
|  | BMI | 0.0376 | 0.0301 |

1. \* Significant after Bonferroni correction.
2. ANI\_Fluency: Animal Fluency; BMI: Body Mass Index; BNT: Boston Naming Task; CVLT\_IM: California Verbal Learning Test Immediate; CVLT\_Long: California Verbal Learning Test Long; DSB: Digit Span Backward; DSF: DSpan igit Forward; DSST: Digital Symbol Substitution Test; MMSE: Mini-Mental State Examination; MOCA: Montreal Cognitive Assessment; TMT\_A: Trail Making Test Part A; TMT\_B: Trail Making Test Part B; VEG\_Fluency: Vegetable Fluency; WM\_SUM: Working Memory Summary.

## Discussion

In this study, we built a brain age prediction model of functional connectivity measures at multiple scales on a large fMRI dataset to learn multiscale functional connectivity patterns of the aging brain and characterize the functional brain. The brain age prediction model built on the harmonized data obtained promising age prediction performance and the derived brain age gap was significantly correlated with cognitive measures, consistent with established brain aging trends.

In the present study, the personalized FNs was computed with an initialization of group-level FNs that were computed in 50 runs, each on a subset of 150 subjects randomly selected from each of the sites with a probability proportional to the sample sizes of different sites. This procedure may generate personalized FNs biased to large sites. We also computed the personalized FNs with an initialization of group-level FNs that were computed in 50 runs, each on a subset of 154 subjects randomly selected from each site with the same number (n = 22) of subjects. Age prediction models were built on the computed personalized FNs and evaluated with the same data harmonization in the tangent space with the same cross-validation. The MAE between the predicted and chronological ages was 5.56 years, close to the result obtained by the prediction models built on the personalized FNs with the group-level FNs computed on subsets with subjects randomly selected proportional to the samples sizes of different sites (MAE = 5.57), indicating that the age prediction performance was not affected too much by the bias in personalized FNs. It merits further investigation to elucidate the sample size effect on the personalized FNs.

Most existing studies investigate the brain functional networks at a specific scale or resolution. For instance, functional networks can be computed based on a set of 17 FNs or a brain atlas with a fixed number of regions. On other hand, recent studies have demonstrated that the brain is a multi-scale system (Betzel and Bassett, 2017; Eickhoff et al., 2018) and with a hierarchical organization (Pines et al., 2022). Therefore, we hypothesized that functional networks computed at multiple scales may provide complementary information for characterizing the brain. Our experiment results have provided empirical evidence that FNs at multiple scales provided useful information for predicting the brain age, with performance better than FNs at any single scale, as indicated by the results presented in Table 2 and Figure 3B. Moreover, the improved brain age prediction performance (Supplementary Table 1) was also observed for functional networks computed with a multiscale brain atlas (Craddock et al., 2012).

We utilized a personalized functional network computational method (Li et al., 2017; Cui et al., 2020). Comparison results have demonstrated that the prediction model built upon the multiscale brain functional connectivity measures could better predict the brain age than those built upon functional connectivity measures of any single scale. As illustrated in Figure 3(A), the multi-scale brain functional networks largely follow a hierarchical organization structure with a few exceptions in finer scales with the FNs computed with settings of 75, 100, 125 and 150 FNs. Specifically, one FN of the LN computed with the setting of 50 FNs switches to the VN computed with the setting of 75 FNs, while two FN of the DMN computed with the setting of 100 FNs switch to SMN and LN computed with the settings of 125 FNs. These results indicated that less stable brain decompositions were generated at finer scales, consistent with findings in structural imaging studies (Varikuti et al., 2018; Patel et al., 2020).

Data harmonization is a prerequisite step in neuroimaging studies with data from multiple sites (Shinohara et al., 2017). Several methods have been developed to harmonize data from different sites, including ComBat and its variants, normative modelling, as well as deep learning based algorithms that transfer the data from different sites into a common, comparable space (Fortin et al., 2017; Fortin et al., 2018; Yu et al., 2018; Dewey et al., 2019; Kia et al., 2020; Moyer et al., 2020; Bayer et al., 2021; Liu et al., 2021; Zuo et al., 2021; Rutherford et al., 2022; Sun et al., 2022). Particularly, ComBat and its variants have been successfully applied to a variety of neuroimaging studies to harmonize diffusion tensor imaging measures, cortical thickness, regional volume measures, and functional connectivity measures (Fortin et al., 2017; Fortin et al., 2018; Yu et al., 2018). A recent study has demonstrated that ComBat-GAM, the method adopted in the present study, outperformed other data harmonization methods in detecting sex differences in regional cortical thickness (Sun et al., 2022). Normative modelling uses percentiles or z-scores to chart the variation of one or several targeting variables orthogonal to the variation of other covariates. Instead of removing site effects from data as a data preprocessing step, normative modelling models site variance as part of the normative model (Kia et al., 2020; Bayer et al., 2021; Rutherford et al., 2022). However, the computational cost and complexity of the model limit its current use to low dimensional imaging data. Deep learning-based data harmonization methods are typically built upon auto-encoders or generative adversarial networks (Dewey et al., 2019; Moyer et al., 2020; Liu et al., 2021; Zuo et al., 2021). Although the deep learning methods have achieved promising performance to translate data from different domains to a common domain, they are limited in their scalability since the deep learning-based data harmonization models have to be trained with objective or loss functions appropriately defined for the problems under study and the training process is computationally heavy. On the other hand, ComBat and its variants are computationally efficient, readily applicable to different data harmonization problems, and capable of modeling both linear and nonlinear effects of covariates. It is out of scope of the present study to compare different data harmonization methods though it is merits further investigation in our future studies.

To improve the generalization ability of machine learning models and increase statistical power, we adopt ComBat-GAM to perform data harmonization. Functional connectivity measures themselves are correlated with each other and reside in a data space that is a manifold, not a vector space (Pervaiz et al., 2020). Therefore, each of the functional connectivity measures should be modeled in conjunction with others for the data harmonization so that potential correlations among different functional connectivity measures can be taken into consideration (Chen et al., 2022). In the present study, we adopt a tangent space modeling method, a successful approach widely adopted in neuroimaging studies (Pervaiz et al., 2020; Zhou et al., 2022), to project the functional connectivity measures onto their tangent space so that the powerful ComBat based data harmonization method can be directly used to harmonize the functional connectivity measures of different sites.

In order to identify informative functional connectivity features for the brain age prediction, we performed permutation tests (Golland et al., 2005) and univariate correlation based significance tests. The statistically significant functional connectivity measures identified by both the tests included 18 connections between SMN, VAN, FPN, and DMN as illustrated in Figure 4. Specifically, many functional network connectivity measures exhibited lower strength with aging, such as those connections with SMN, DMN, and FPN, consistent with findings of the existing studies (Chan et al., 2014; Grady et al., 2016; Zonneveld et al., 2019; Truelove-Hill et al., 2020). Interestingly, the within network connectivity strength of the VN and SMN became higher with aging. Similar findings were also reported in several existing studies (Song et al., 2014; Seidler et al., 2015; Zonneveld et al., 2019). We noticed that the connectivity between DMN (network 16, the anterior cingulate) and VAN (network 1, the insula) computed with the setting of 17 FNs exhibited an increasing pattern with aging, consistent with findings reported in previous studies (Cao et al., 2014; Fan et al., 2020), as well as higher connectivity between SMN and DAN computed with the settings of 17 and 50 FNs. Similar to findings reported in (de Lange et al., 2020), we also found that the most informative FC measures for the brain age prediction might vary across the lifespan, as shown in Figure S1 of the supplementary data.

A highly consistent finding in the aging literature is lower DMN connectivity with aging (Ferreira and Busatto, 2013; Dennis and Thompson, 2014; Damoiseaux, 2017; Stumme et al., 2020). Aging is also associated with substantial declines in motor functioning as well as higher-order cognitive networks, such as the FPN. In addition to these findings, we also observed that aging was associated with higher within-network connectivity in the VN and SMN, contradictory with findings of existing studies (Stumme et al., 2020). Particularly, it was observed that aging is associated with lower functional connectivity within the primary processing networks, including the VN and SMN (Stumme et al., 2020). A variety of factors might contribute to such discrepancies, including different atlases or brain parcellations used for computing the functional connectivity measure (Arslan et al., 2018) and the sample size and diversity (Marek et al., 2022).

Many methods are available to evaluate feature importance and interpretability of machine learning models (Lundberg and Lee, 2017; Hou and Zhou, 2020), including the label permutation method adopted in the present study and many model-agnostic methods, such as permutation feature importance, partial dependence plot, and Shapely values. We chose the label permutation method because it is capable of generating statistical significance for feature importance and taking into consideration of interactive effects of all features. We will explore other feature importance evaluation methods in our future studies since all the methods have their specific advantages and limitations.

Recent studies have demonstrated that the BAG score could potentially serve as a quantitative marker of the brain aging (Cole et al., 2017; Smith et al., 2019; Boyle et al., 2021; Dinsdale et al., 2021). In order to investigate to what extent the BAG score is associated with cognitive and clinical measures of the subjects in the multisite cohort, Pearson correlation analysis was performed between the BAG score and each of the available cognitive and clinical measures, including DSF (Digit Span Forward), DSST (Digital Symbol Substitution Test), DSB (Digit Span Backward), MMSE (Mini-Mental State Examination), MOCA (Montreal Cognitive Assessment), WM_SUM (Working Memory Summary), verbal fluency and verbal memory related tasks, TMT_A and TMT_B (Trail Making Test Part A and B), Systole, Diastole, and BMI (Body Mass Index). In all the analyses, age, sex, and site were included as covariates. We found that the BAG score was positively correlated with measures decreasing with aging and negatively correlated with those increasing with aging, consisting with existing findings that a positive BAG score indicated an accelerated aging process (Cole et al., 2017; Smith et al., 2019; Boyle et al., 2021; Dinsdale et al., 2021).

Our study has following limitations. First, the present study has focused on functional connectivity within FNs. Functional connectivity within networks across different scales may provide additional informative functional connectivity measures for characterizing the aging brain (Iraji et al., 2021). Second, the present study has focused on functional connectivity information alone. The functional connectivity measures can be further enhanced by functional network topology measures that are informative for charactering the brain development (Cui et al., 2020). Moreover, integration of brain anatomy, structural connectivity, and functional connectivity measures may further improve the brain age prediction as well as the characterization of brain aging (Truelove-Hill et al., 2020). Third, the present study identified informative functional connectivity measures using a combination of permutation test and univariate correlation analysis. It merits further investigation to establish explainable and interpretable brain age prediction models (Tian and Zalesky, 2021).

In summary, the present study revealed that functional connectivity measures at multiple scales were more informative than those at any single spatial scale for the brain age prediction and the data harmonization in the tangent space of functional connectivity measures significantly improved the brain age prediction performance. Moreover, the derived brain age gap score was associated with cognitive and clinical measures.

## Competing Financial Interests

Dr. Nasrallah was an educational speaker for Biogen. Dr. Wolk has received grant support from Merck, Biogen, and Eli Lilly/Avid and consultation fees from Neuronix and GE Healthcare and is on the Data and Safety Monitoring Board for a clinical trial run by Functional Neuromodulation.

## Acknowledgement

This work was partially supported, in part, by NIH grants (RF1-AG054409, U01-AG068057, R01-AG066650, R01-EB022573 and R01-MH123550). The BLSA is supported by the Intramural Research Program, National Institute of Aging, and Research and Development Contract HHSN-260-2004-00012C.

## Credit authorship contribution statement

**Zhen Zhou**: Conceptualization, methodology, coding, validation, writing – original draft, writing – review & editing, visualization. **Dhivya Srinivasan**: Data analysis. **Hongming Li**: Conceptualization, methodology, coding, writing – review. **Ahmed Abdulkadir**: Conceptualization, methodology, writing – review. **Ilya Nasrallah**: Writing – review. **Junhao Wen**: Writing – review. **Jimit Doshi**: Data analysis, writing – review. **Guray Erus**: Data analysis, writing – review. **Elizabeth Mamourian**: Data analysis. **Nick R. Bryan**: Writing – review. **David A. Wolk**: Writing – review. **Lori Beason-Held**: Writing – review. **Susan M. Resnick**: Writing – review. **Theodore D. Satterthwaite**: Writing – review. **Christos Davatzikos**: Conceptualization, methodology, writing – review. **Haochang Shou**: Conceptualization, methodology, writing – review. **Yong Fan**: Conceptualization, methodology, writing – revision.

## Data and code availability

The code to compute personalized functional network is available at https://github.com/hmlicas/Collaborative_Brain_Decomposition. Combat data harmonization was performed using a package available at https://github.com/rpomponio/neuroHarmonize. The data of this study is available upon request and subject to approval by the ISTAGING Consortium’s supervisory committee. All the other codes will be deposited in an open access platform upon the acceptance of this manuscript.

## Notes

### Summary of Updates

More detailed are added in Materials and Methods; Discussion updated; Supplemental files added.

## References

Alfaro-Almagro F et al. (2018) Image processing and Quality Control for the first 10,000 brain imaging datasets from UK Biobank. Neuroimage 166:400–424.

Arslan S, Ktena SI, Makropoulos A, Robinson EC, Rueckert D, Parisot S (2018) Human brain mapping: A systematic comparison of parcellation methods for the human cerebral cortex. Neuroimage 170:5–30.

Bashyam VM et al. (2020) MRI signatures of brain age and disease over the lifespan based on a deep brain network and 14 468 individuals worldwide. Brain 143:2312–2324.

Bayer JMM, Dinga R, Kia SM, Kottaram AR, Wolfers T, Lv J, Zalesky A, Schmaal L, Marquand A (2021) Accommodating site variation in neuroimaging data using normative and hierarchical Bayesian models.

Beheshti I, Nugent S, Potvin O, Duchesne S (2019) Bias-adjustment in neuroimaging-based brain age frameworks: A robust scheme. NeuroImage: Clinical 24:102063.

Betzel RF, Bassett DS (2017) Multi-scale brain networks. Neuroimage 160:73–83.

Boyle R, Jollans L, Rueda-Delgado LM, Rizzo R, Yener GG, McMorrow JP, Knight SP, Carey D, Robertson IH, Emek-Savas DD, Stern Y, Kenny RA, Whelan R (2021) Brain-predicted age difference score is related to specific cognitive functions: a multi-site replication analysis. Brain Imaging Behav 15:327–345.

Cao W, Luo C, Zhu B, Zhang D, Dong L, Gong J, Gong D, He H, Tu S, Yin W, Li J, Chen H, Yao D (2014) Resting-state functional connectivity in anterior cingulate cortex in normal aging. Front Aging Neurosci 6:280.

Chan MY, Park DC, Savalia NK, Petersen SE, Wig GS (2014) Decreased segregation of brain systems across the healthy adult lifespan. Proc Natl Acad Sci U S A 111:E4997–5006.

Chen AA, Srinivasan D, Pomponio R, Fan Y, Nasrallah IM, Resnick SM, Beason-Held LL, Davatzikos C, Satterthwaite TD, Bassett DS, Shinohara RT, Shou H (2022) Harmonizing functional connectivity reduces scanner effects in community detection. Neuroimage 256:119198.

Chen J, Liu J, Calhoun VD, Arias-Vasquez A, Zwiers MP, Gupta CN, Franke B, Turner JA (2014) Exploration of scanning effects in multi-site structural MRI studies. J Neurosci Methods 230:37–50.

Chung HK, Tymula A, Glimcher P (2017) The Reduction of Ventrolateral Prefrontal Cortex Gray Matter Volume Correlates with Loss of Economic Rationality in Aging. J Neurosci 37:12068–12077.

Cole JH, Franke K (2017) Predicting Age Using Neuroimaging: Innovative Brain Ageing Biomarkers. Trends Neurosci 40:681–690.

Cole JH, Poudel RPK, Tsagkrasoulis D, Caan MWA, Steves C, Spector TD, Montana G (2017) Predicting brain age with deep learning from raw imaging data results in a reliable and heritable biomarker. Neuroimage 163:115–124.

Craddock RC, James GA, Holtzheimer PE, 3rd, Hu XP, Mayberg HS (2012) A whole brain fMRI atlas generated via spatially constrained spectral clustering. Hum Brain Mapp 33:1914–1928.

Cui Z, Su M, Li L, Shu H, Gong G (2018) Individualized Prediction of Reading Comprehension Ability Using Gray Matter Volume. Cereb Cortex 28:1656–1672.

Cui Z et al. (2020) Individual Variation in Functional Topography of Association Networks in Youth. Neuron 106:340–353 e348.

Cui Z et al. (2022) Linking Individual Differences in Personalized Functional Network Topography to Psychopathology in Youth. Biological Psychiatry.

Damoiseaux JS (2017) Effects of aging on functional and structural brain connectivity. Neuroimage 160:32–40.

de Lange AG, Anaturk M, Suri S, Kaufmann T, Cole JH, Griffanti L, Zsoldos E, Jensen DEA, Filippini N, Singh-Manoux A, Kivimaki M, Westlye LT, Ebmeier KP (2020) Multimodal brain-age prediction and cardiovascular risk: The Whitehall II MRI sub-study. Neuroimage 222:117292.

Dennis EL, Thompson PM (2014) Functional brain connectivity using fMRI in aging and Alzheimer’s disease. Neuropsychol Rev 24:49–62.

Dewey BE, Zhao C, Reinhold JC, Carass A, Fitzgerald KC, Sotirchos ES, Saidha S, Oh J, Pham DL, Calabresi PA, van Zijl PCM, Prince JL (2019) DeepHarmony: A deep learning approach to contrast harmonization across scanner changes. Magn Reson Imaging 64:160–170.

Di Martino A, Fair DA, Kelly C, Satterthwaite TD, Castellanos FX, Thomason ME, Craddock RC, Luna B, Leventhal BL, Zuo XN, Milham MP (2014) Unraveling the miswired connectome: a developmental perspective. Neuron 83:1335–1353.

Dinsdale NK, Bluemke E, Smith SM, Arya Z, Vidaurre D, Jenkinson M, Namburete AIL (2021) Learning patterns of the ageing brain in MRI using deep convolutional networks. Neuroimage 224:117401.

Dosenbach NU, Nardos B, Cohen AL, Fair DA, Power JD, Church JA, Nelson SM, Wig GS, Vogel AC, Lessov-Schlaggar CN, Barnes KA, Dubis JW, Feczko E, Coalson RS, Pruett JR, Jr., Barch DM, Petersen SE, Schlaggar BL (2010) Prediction of individual brain maturity using fMRI. Science 329:1358–1361.

Douaud G, Groves AR, Tamnes CK, Westlye LT, Duff EP, Engvig A, Walhovd KB, James A, Gass A, Monsch AU, Matthews PM, Fjell AM, Smith SM, Johansen-Berg H (2014) A common brain network links development, aging, and vulnerability to disease. Proc Natl Acad Sci U S A 111:17648–17653.

Eickhoff SB, Yeo BTT, Genon S (2018) Imaging-based parcellations of the human brain. Nat Rev Neurosci 19:672–686.

Erus G, Battapady H, Satterthwaite TD, Hakonarson H, Gur RE, Davatzikos C, Gur RC (2015) Imaging patterns of brain development and their relationship to cognition. Cereb Cortex 25:1676–1684.

Fair DA, Cohen AL, Dosenbach NU, Church JA, Miezin FM, Barch DM, Raichle ME, Petersen SE, Schlaggar BL (2008) The maturing architecture of the brain’s default network. Proc Natl Acad Sci U S A 105:4028–4032.

Fan X, Li H, Li K (2020) Increased functional connectivity of anterior insula to anterior cingulate cortex in amnestic mild cognitive impairment: A longitudinal restingl_state fMRI study. Alzheimer’s & Dementia 16.

Ferreira LK, Busatto GF (2013) Resting-state functional connectivity in normal brain aging. Neurosci Biobehav Rev 37:384–400.

Filippini N et al. (2014) Study protocol: the Whitehall II imaging sub-study. BMC Psychiatry 14:159.

Fletcher PT, Lu C, Pizer SM, Joshi S (2004) Principal geodesic analysis for the study of nonlinear statistics of shape. IEEE Trans Med Imaging 23:995–1005.

Focke NK, Helms G, Kaspar S, Diederich C, Toth V, Dechent P, Mohr A, Paulus W (2011) Multisite voxel-based morphometry--not quite there yet. Neuroimage 56:1164–1170.

Fortin J-P, Parker D, Tunç B, Watanabe T, Elliott MA, Ruparel K, Roalf DR, Satterthwaite TD, Gur RC, Gur RE (2017) Harmonization of multi-site diffusion tensor imaging data. Neuroimage 161:149–170.

Fortin JP, Cullen N, Sheline YI, Taylor WD, Aselcioglu I, Cook PA, Adams P, Cooper C, Fava M, McGrath PJ, McInnis M, Phillips ML, Trivedi MH, Weissman MM, Shinohara RT (2018) Harmonization of cortical thickness measurements across scanners and sites. Neuroimage 167:104–120.

Golland P, Liang F, Mukherjee S, Panchenko D (2005) Permutation Tests for Classification. In, pp 501–515. Berlin, Heidelberg: Springer Berlin Heidelberg.

Grady C, Sarraf S, Saverino C, Campbell K (2016) Age differences in the functional interactions among the default, frontoparietal control, and dorsal attention networks. Neurobiol Aging 41:159–172.

Griffanti L, Salimi-Khorshidi G, Beckmann CF, Auerbach EJ, Douaud G, Sexton CE, Zsoldos E, Ebmeier KP, Filippini N, Mackay CE, Moeller S, Xu J, Yacoub E, Baselli G, Ugurbil K, Miller KL, Smith SM (2014) ICA-based artefact removal and accelerated fMRI acquisition for improved resting state network imaging. Neuroimage 95:232–247.

Habes M, Erus G, Toledo JB, Zhang T, Bryan N, Launer LJ, Rosseel Y, Janowitz D, Doshi J, Van der Auwera S, von Sarnowski B, Hegenscheid K, Hosten N, Homuth G, Volzke H, Schminke U, Hoffmann W, Grabe HJ, Davatzikos C (2016) White matter hyperintensities and imaging patterns of brain ageing in the general population. Brain 139:1164–1179.

Habes M et al. (2021) The Brain Chart of Aging: Machine-learning analytics reveals links between brain aging, white matter disease, amyloid burden, and cognition in the iSTAGING consortium of 10,216 harmonized MR scans. Alzheimers Dement 17:89–102.

Hou BJ, Zhou ZH (2020) Learning With Interpretable Structure From Gated RNN. IEEE Trans Neural Netw Learn Syst 31:2267–2279.

Iraji A, Faghiri A, Fu Z, Rachakonda S, Kochunov P, Belger A, Ford JM, McEwen S, Mathalon DH, Mueller BA, Pearlson GD, Potkin SG, Preda A, Turner JA, van Erp TGM, Calhoun VD (2021) Multi-spatial scale dynamic interactions between functional sources reveal sex-specific changes in schizophrenia. Network Neuroscience:1–48.

Jenkinson M, Beckmann CF, Behrens TE, Woolrich MW, Smith SM (2012) Fsl. Neuroimage 62:782–790.

Kaufmann T et al. (2019) Common brain disorders are associated with heritable patterns of apparent aging of the brain. Nat Neurosci 22:1617–1623.

Kia SM, Huijsdens H, Dinga R, Wolfers T, Mennes M, Andreassen OA, Westlye LT, Beckmann CF, Marquand AF (2020) Hierarchical Bayesian Regression for Multi-site Normative Modeling of Neuroimaging Data. In: Medical Image Computing and Computer Assisted Intervention – MICCAI 2020, pp 699–709.

Li H, Satterthwaite TD, Fan Y (2017) Large-scale sparse functional networks from resting state fMRI. Neuroimage 156:1–13.

Li H, Satterthwaite TD, Fan Y (2018) Brain age prediction based on resting-state functional connectivity patterns using convolutional neural networks. In: 2018 IEEE 15th International Symposium on Biomedical Imaging (ISBI 2018), pp 101–104.

Liang H, Zhang F, Niu X (2019) Investigating systematic bias in brain age estimation with application to post-traumatic stress disorders. Hum Brain Mapp 40:3143–3152.

Liu M, Maiti P, Thomopoulos S, Zhu A, Chai Y, Kim H, Jahanshad N (2021) Style Transfer Using Generative Adversarial Networks for Multi-Site MRI Harmonization. Med Image Comput Comput Assist Interv 12903:313–322.

Lundberg SM, Lee S-I (2017) A unified approach to interpreting model predictions. Advances in neural information processing systems 30.

Marek S et al. (2022) Reproducible brain-wide association studies require thousands of individuals. Nature 603:654–660.

Minkova L, Habich A, Peter J, Kaller CP, Eickhoff SB, Kloppel S (2017) Gray matter asymmetries in aging and neurodegeneration: A review and meta-analysis. Hum Brain Mapp 38:5890–5904.

Mourao-Miranda J, Bokde AL, Born C, Hampel H, Stetter M (2005) Classifying brain states and determining the discriminating activation patterns: Support Vector Machine on functional MRI data. Neuroimage 28:980–995.

Moyer D, Ver Steeg G, Tax CMW, Thompson PM (2020) Scanner invariant representations for diffusion MRI harmonization. Magn Reson Med 84:2174–2189.

Ng B, Dressler M, Varoquaux G, Poline JB, Greicius M, Thirion B (2014) Transport on Riemannian manifold for functional connectivity-based classification. Med Image Comput Comput Assist Interv 17:405–412.

Patel R, Steele CJ, Chen AGX, Patel S, Devenyi GA, Germann J, Tardif CL, Chakravarty MM (2020) Investigating microstructural variation in the human hippocampus using non-negative matrix factorization. Neuroimage 207:116348.

Pervaiz U, Vidaurre D, Woolrich MW, Smith SM (2020) Optimising network modelling methods for fMRI. Neuroimage 211:116604.

Pines AR et al. (2022) Dissociable multi-scale patterns of development in personalized brain networks. Nat Commun 13:2647.

Pomponio R et al. (2020) Harmonization of large MRI datasets for the analysis of brain imaging patterns throughout the lifespan. Neuroimage 208:116450.

Prins ND, Scheltens P (2015) White matter hyperintensities, cognitive impairment and dementia: an update. Nat Rev Neurol 11:157–165.

Rutherford S, Kia SM, Wolfers T, Fraza C, Zabihi M, Dinga R, Berthet P, Worker A, Verdi S, Ruhe HG, Beckmann CF, Marquand AF (2022) The normative modeling framework for computational psychiatry. Nat Protoc 17:1711–1734.

Salimi-Khorshidi G, Douaud G, Beckmann CF, Glasser MF, Griffanti L, Smith SM (2014) Automatic denoising of functional MRI data: combining independent component analysis and hierarchical fusion of classifiers. Neuroimage 90:449–468.

Seidler R, Erdeniz B, Koppelmans V, Hirsiger S, Merillat S, Jancke L (2015) Associations between age, motor function, and resting state sensorimotor network connectivity in healthy older adults. Neuroimage 108:47–59.

Shanmugan S et al. (2022) Sex differences in the functional topography of association networks in youth. Proc Natl Acad Sci U S A 119:e2110416119.

Shinohara RT et al. (2017) Volumetric Analysis from a Harmonized Multisite Brain MRI Study of a Single Subject with Multiple Sclerosis. AJNR Am J Neuroradiol 38:1501–1509.

Smith SM, Vidaurre D, Alfaro-Almagro F, Nichols TE, Miller KL (2019) Estimation of brain age delta from brain imaging. Neuroimage 200:528–539.

Song J, Birn RM, Boly M, Meier TB, Nair VA, Meyerand ME, Prabhakaran V (2014) Age-related reorganizational changes in modularity and functional connectivity of human brain networks. Brain Connect 4:662–676.

Stumme J, Jockwitz C, Hoffstaedter F, Amunts K, Caspers S (2020) Functional network reorganization in older adults: Graph-theoretical analyses of age, cognition and sex. Neuroimage 214:116756.

Sun D et al. (2022) A comparison of methods to harmonize cortical thickness measurements across scanners and sites. Neuroimage 261:119509.

Tian Y, Zalesky A (2021) Machine learning prediction of cognition from functional connectivity: Are feature weights reliable? NeuroImage.

Truelove-Hill M, Erus G, Bashyam V, Varol E, Sako C, Gur RC, Gur RE, Koutsouleris N, Zhuo C, Fan Y, Wolf DH, Satterthwaite TD, Davatzikos C (2020) A Multidimensional Neural Maturation Index Reveals Reproducible Developmental Patterns in Children and Adolescents. J Neurosci 40:1265–1275.

Varikuti DP, Genon S, Sotiras A, Schwender H, Hoffstaedter F, Patil KR, Jockwitz C, Caspers S, Moebus S, Amunts K, Davatzikos C, Eickhoff SB (2018) Evaluation of non-negative matrix factorization of grey matter in age prediction. Neuroimage 173:394–410.

Yger F, Berar M, Lotte F (2017) Riemannian Approaches in Brain-Computer Interfaces: A Review. IEEE Trans Neural Syst Rehabil Eng 25:1753–1762.

You K, Park HJ (2021) Re-visiting Riemannian geometry of symmetric positive definite matrices for the analysis of functional connectivity. Neuroimage 225:117464.

Yu M, Linn KA, Cook PA, Phillips ML, McInnis M, Fava M, Trivedi MH, Weissman MM, Shinohara RT, Sheline YI (2018) Statistical harmonization corrects site effects in functional connectivity measurements from multi-site fMRI data. Hum Brain Mapp 39:4213–4227.

Zhou Z, Srinivasan D, Li H, Abdulkadir A, Shou H, Davatzikos C, Fan Y, Gimi BS, Krol A (2022) Harmonization of multi-site functional connectivity measures in tangent space improves brain age prediction. In: Medical Imaging 2022: Biomedical Applications in Molecular, Structural, and Functional Imaging.

Zhou Z, Chen XB, Zhang Y, Hu D, Qiao LS, Yu RP, Yap PT, Pan G, Zhang H, Shen DG (2020) A toolbox for brain network construction and classification (BrainNetClass). Human Brain Mapping 41:2808–2826.

Zonneveld HI, Pruim RH, Bos D, Vrooman HA, Muetzel RL, Hofman A, Rombouts SA, van der Lugt A, Niessen WJ, Ikram MA, Vernooij MW (2019) Patterns of functional connectivity in an aging population: The Rotterdam Study. Neuroimage 189:432–444.

Zuo L, Dewey BE, Liu Y, He Y, Newsome SD, Mowry EM, Resnick SM, Prince JL, Carass A (2021) Unsupervised MR harmonization by learning disentangled representations using information bottleneck theory. Neuroimage 243:118569.

