## Supplemental Materials for "Multiscale functional connectivity patterns of the aging brain learned from rsfMRI data of 4,259 individuals of the multi-cohort iSTAGING study"

We have also evaluated brain age prediction models built on FC measures computed based on Craddock’s multiscale atlases, generated by 2-level group clustering using temporal correlation (available at <http://ccraddock.github.io/cluster_roi/>) at scale 20, 30, 50, 70, 100, 120, and 150 (Craddock et al., 2012). The multiscale FC measures were first harmonized measures in the tangent space) and then were used to build brain age prediction models that were evaluated with the same procedure of the present study. As summarized in Table S1, the best performance was obtained by the model build on the multiscale FC measures. These results indicated that multiscale FC measures could achieve better brain age prediction performance when a different method was used to compute the FC measures.

Table S1. Comparison of age prediction performance using different brain parcellations

| Personalized network | Scale | 17 | 25 | 50 | 75 | 100 | 125 | 150 | Multiscale |
| --- | --- | --- | --- | --- | --- | --- | --- | --- | --- |
|  | MAE | 7.96 | 7.43 | 7.33 | 8.17 | 7.80 | 6.96 | 6.45 | 5.57 |
| Craddock  atlas | Scale | 20 | 30 | 50 | 70 | 100 | 120 | 150 | Multiscale |
|  | MAE | 7.83 | 7.33 | 7.30 | 8.30 | 8.24 | 6.85 | 6.14 | 5.70 |

We compared brain age prediction models built using different regression methods, including support vector regression (SVR) and XGBoost on the multiscale FC measures. Particularly, we adopted the same nested 5-fold cross-validation for evaluating all the three methods, with the inner cross-validation to determine the optimal parameters. For SVR, C was tuned within the set of [1, 2^2^, 2^3^, 2^4^, 2^5^]; For ridge regression, λ was tuned within the set of [2^3^, 2^3.5^, 2^4^, …, 2^7.5^, 2^8^]. As grid search for multiple parameters could be difficult, we applied a random search for the parameters in XGBoost (Chen and Guestrin, 2016): ‘min_child_weight’ in [1, 2, …, 6], ‘reg_lambda’ in a log-uniform distribution from 10 to 100, ‘subsample’ in a uniform distribution from 0.5 to 1, ‘colsample_bytree’ in a uniform distribution from 0.5 to 1, ‘n_estimators’ in [200, …, 500] and ‘max_depth’ in [2, 3, 4, 5]. The number of random search iterations was set to 100. As demonstrated in the Table S2, ridge regression outperformed the other two methods in terms of both computation time and model performance.

Table S2 Comparison of computation time and performance of different predictive methods

| MAE (year) between predicted age and chronological age | | | Computation time | | |
| --- | --- | --- | --- | --- | --- |
| SVR | XGBoost | Ridge | SVR | XGBoost | Ridge |
| 6.28 | 5.70 | 5.57 | 32 hours 9 mins | 187 hours 12 mins | 4 mins 19 s |

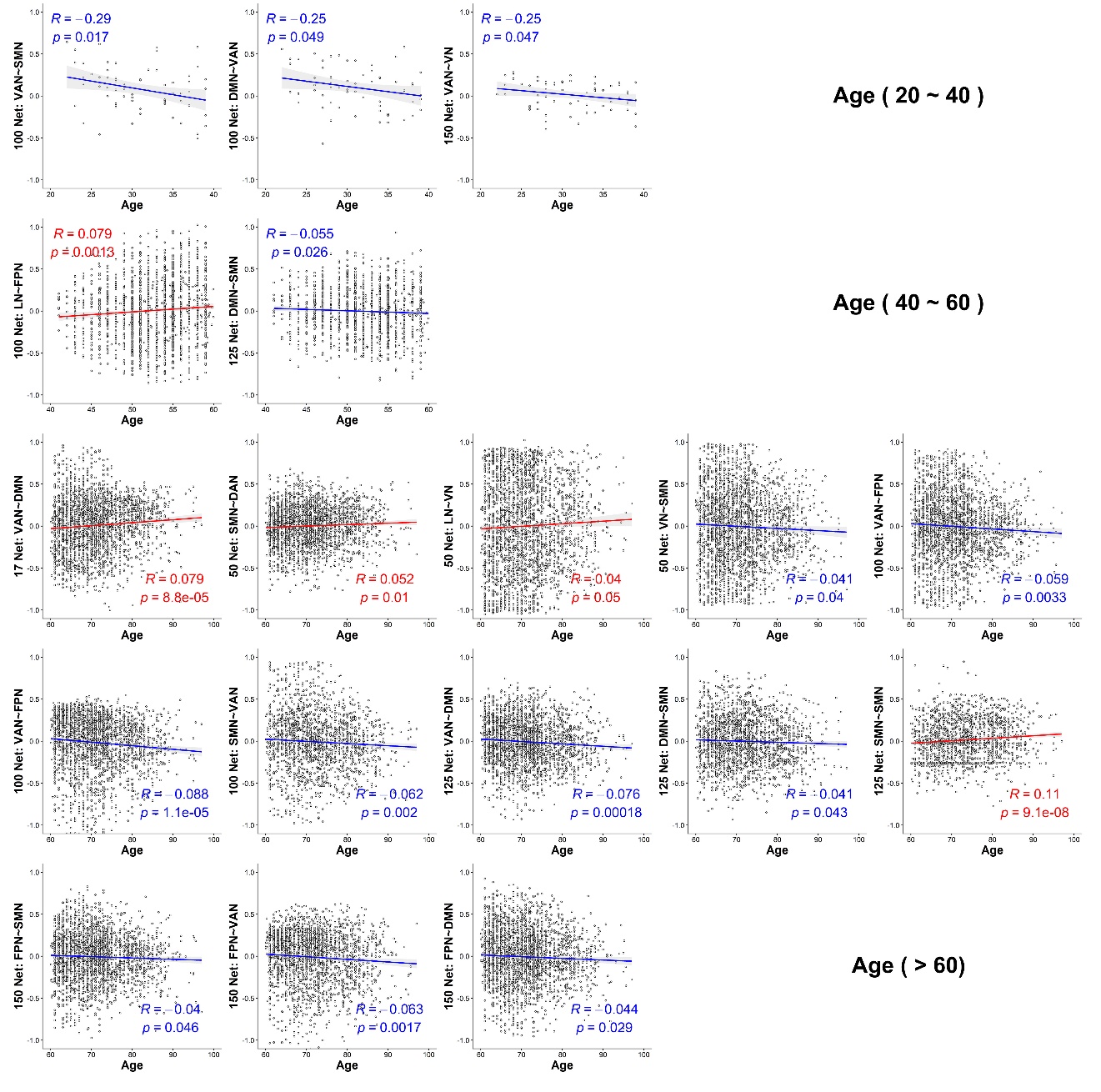

**Figure S1.** Functional connections with connectivity significantly correlated with the chronological age in three age ranges (20 ~ 40 years, 40 ~ 60 years, and > 60 years).

We explored whether the FC measures informative for the age prediction varied across age groups. Specifically, we computed correlations between age and FC measures that were informative for the age prediction for subjects in three different age groups, including younger than 40 years( n=65), between 40 and 60 years (n = 1641), and older than 60 years (n = 2461) subjects older than 60). As shown in Figure r1, three FC measures were significantly correlated with age in the group younger than 40 years, two FC measures were significantly correlated with age in the group between 40 and 60 years old, and 13 FC measures were significantly correlated with age in the group older than 60 years, different from those shown in Figure 4. These results indicated that the most informative features for the brain age prediction might vary across the lifespan, with a caveat that the results were obtained based on various age groups with different sample sizes.
